# Comparison of unit resolution versus high-resolution accurate mass for parallel reaction monitoring

**DOI:** 10.1101/2021.05.04.442680

**Authors:** Lilian R. Heil, Philip M. Remes, Michael J. MacCoss

## Abstract

Parallel reaction monitoring (PRM) is an increasingly popular alternative to selected reaction monitoring (SRM) for targeted proteomics. PRM’s strengths over SRM are that it monitors all product ions in a single spectrum, thus eliminating the need to select interference-free product ions prior to data acquisition, and that it is most frequently performed on high-resolution instruments, such as quadrupole-orbitrap and quadrupole-time of flight instruments. Here, we show that the primary advantage of PRM is the ability to monitor all transitions in parallel, and that high-resolution data are not necessary to obtain high quality quantitative data. We run the same scheduled PRM assay, measuring 432 peptides from 126 plasma proteins, multiple times on a Orbitrap Eclipse Tribrid mass spectrometer, alternating separate liquid chromatography-tandem mass spectrometry runs between the high resolution Orbitrap and the unit resolution linear ion trap for PRM. We find that both mass analyzers have similar technical precision, and that the linear ion trap’s superior sensitivity gives it better lower limits of quantitation on over 62% of peptides in the assay.

**Abstract graphic:** 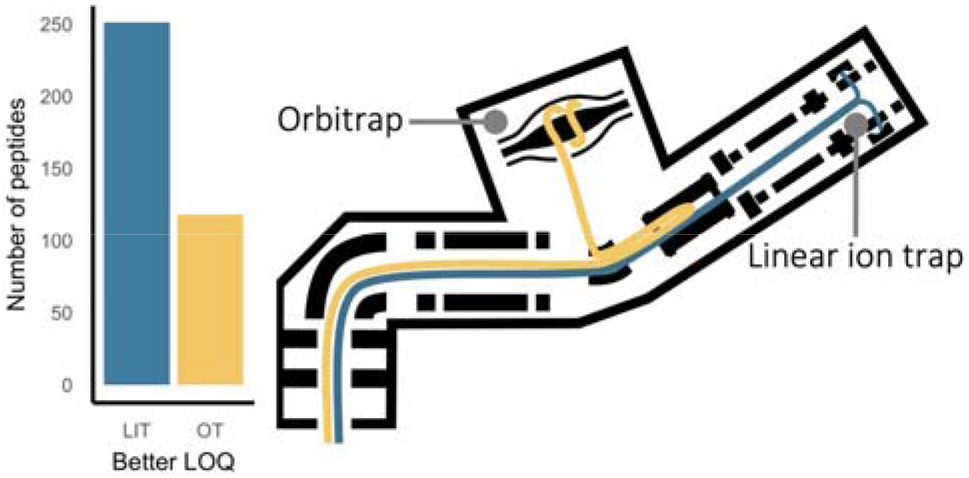

## Introduction

Targeted proteomics is a strategy to obtain accurate and precise quantitation by maximizing the time a mass spectrometer (MS) spends measuring a pre-defined subset of analytes in a sample. Historically, targeted assays have been performed using selected reaction monitoring (SRM) on a triple quadrupole mass spectrometer.^1^ In an SRM assay, a specific list of precursor and product ion pairs, called transitions, are monitored by setting the quadrupoles to transmit specific reactions at specific times during the liquid chromatography (LC) gradient.^2,3^ SRM is widely used today due to its reliability and versatility, but it still has limitations. Notably, the need to select interference-free transitions can limit the selectivity and breadth of an experiment; because each precursor/ product ion pair needs to be targeted individually, each additional product ion measured comes at a cost to overall cycle time. Consequently, there is a tradeoff between increasing selectivity and measuring more precursor ions because targeting more precursors means sacrificing selectivity by monitoring fewer transitions per precursor. Additionally, any change to the set of product ions requires rerunning the entire experiment.

Parallel reaction monitoring (PRM) is an alternative to SRM where the entire product ion spectrum is collected for a target precursor instead of selected precursor-product ion reactions. Early versions of PRM showed the potential to benefit from the full scanning capability of a unit-resolution ion trap.^4^ However, PRM was popularized on high resolution quadrupole-Orbitrap (q-OT)^5,6^ and quadrupole-Time of Flight^7^ mass spectrometers. Since these demonstrations, the performance of PRM was largely attributed to two factors: the ability to monitor more product ions per precursor and the higher confidence in product ions associated with the high resolution product ion spectra.^5,8,9^ However, we hypothesize that the primary benefit of PRM is the ability to measure many product ions in a single scan, and that high resolution spectra are not critical to its success.

To assess the importance of high resolution mass analyzers in PRM, we sought to compare the performance of a unit resolution quadrupole linear ion trap (q-LIT) to a high-resolution q-OT for PRM experiments. We use an Orbitrap Eclipse Tribrid instrument to run a controlled experiment and establish that a q-LIT can be used on its own for targeted quantitative proteomics assays without sacrificing significant precision or quantitative accuracy compared to the q-OT. This instrument has both an Orbitrap and a linear ion trap that can be operated independently which allows us to inject the same sample multiple times, alternating between the Orbitrap and linear ion trap with each injection, thereby eliminating the variability inherent to comparisons where multiple instruments are used (**Figure 1**).

**Figure 1.**
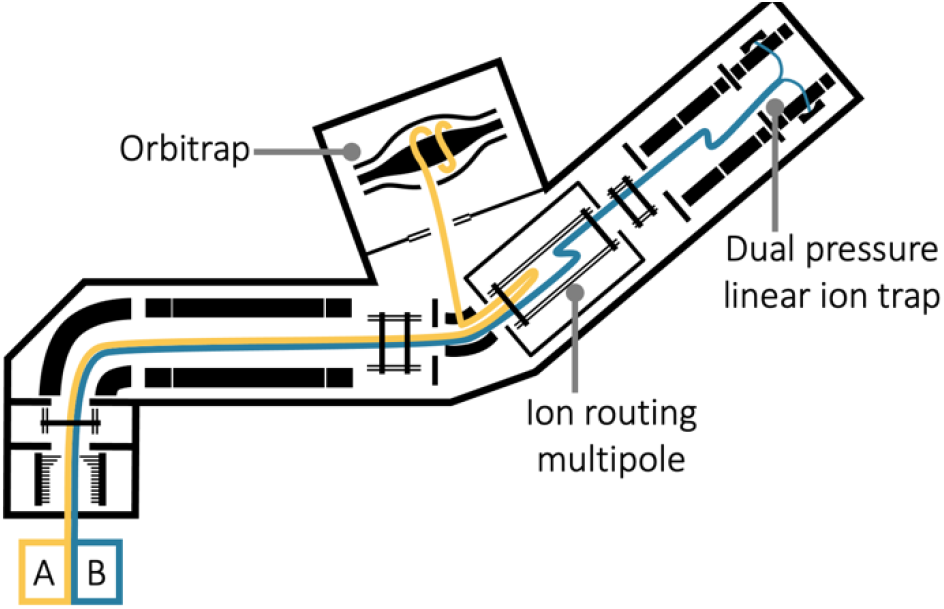
Schematic of the Orbitrap Fusion Eclipse Tribrid instrument. Ion paths are highlighted in yellow and blue. To compare Orbitrap to ion trap directly, alternate injections were performed using each mass analyzer. Injections either followed path A, where the instrument operated as a quadrupole-Orbitrap or path B, where the instrument operated as a quadrupole-linear ion trap.

## Experimental

### Sample preparation and mass spectrometry

#### Sample preparation

A 50:50 mix of female and male pooled human plasma from Golden West Biologicals was created (Lot # MSG11500F and MSG11500M). Both the pooled human plasma and chicken serum (Cat #SC30 Lot #190509-0130, Equitech-Bio, Inc.) were split into aliquots containing approximately 300 *μ*g of total protein. Both aliquots were reduced, alkylated, and digested with trypsin using S-Trap Mini sample preparation kit (ProtiFi) according to manufacturer guidelines.^10^ The digested samples were diluted to a concentration of approximately 0.1 mg/mL with 0.1% formic acid in water with Pierce Peptide Retention Time Standard (PRTC) (Thermo Fisher Scientific) at a concentration of 0.4 nmol/mL. A dilution series was prepared by diluting the human plasma digest in the chicken serum digest at 11 different dilution points of 0, 0.5, 1, 3, 5, 7, 10, 30, 50, 70 and 100% human plasma (**Table S1**).

#### Liquid Chromatography-Mass Spectrometry

LC-MS/MS analysis was performed with a Thermo Easy-nanoLC coupled to a Thermo Orbitrap Eclipse Tribrid Mass spectrometer. Peptides were separated using an in-house manufactured column created by pulling 75 *μ*m inner diameter fused silica capillary (TSP075375, Molex) to an approximately 5 *μ*m ID tip using a Sutter Instruments CO2 laser based micropipette puller. The pulled tip capillary was packed with 30 cm of 3 *μ*m ReproSil-Pur C18 beads (Dr. Maisch GmbH) using an in-house developed pressure bomb with high pressure helium gas. The trap column consisted 150 *μ*m inner diameter fused silica capillary (TSP150375, Molex) fritted on one end with Kasil and packed with 4 cm of 3 *μ*m ReproSil-Pur C18 beads (Dr. Maisch GmbH). Mobile-phase A was 0.1% formic acid in water and mobile-phase B was 0.1% formic acid in 80% acetonitrile. Peptides were separated with reversed-phase liquid chromatography over a 90 minute gradient from 0 to 40% B followed by a 5 minute ramp to 75% B, a 5 minute hold at B, and an additional 15 minute wash at 100% B.

Data for the chromatogram library were acquired previously using the same chromatography setup coupled to a Thermo Lumos Tribrid Mass spectrometer. Tandem mass spectrometry data were acquired in 6 gas-phase fractions with 4-*m/z* staggered data independent acquisition (DIA) windows as described in detail by Pino et al. (2020).^11^ Retention time calibration was performed on the Eclipse with the same LC method using 8-*m/z* staggered DIA window isolation scheme described previously.^11^

#### Parallel Reaction monitoring

All parallel reaction monitoring experiments were performed using one of four methods. The four methods can be distinguished based on the following characteristics (analyzer, quadrupole precursor isolation width): linear ion trap, 0.7 Th; linear ion trap, 1.7 Th; Orbitrap, 0.7 Th; and Orbitrap, 1.7 Th. Each method featured a standard MS1 scan and a set of targeted MS2 scans, all acquired with the same mass analyzer. The same target list with scheduled acquisition was used in all methods, and scheduling was adjusted throughout the experiment to account for retention time drift. For MS2 acquisition, all methods used standard AGC target, dynamic maximum inject time with 20 desired points across the expected peak width of 40 s. Fragmentation occurred in the ion routing multipole using HCD with normalized collision energy of 30%. When the linear ion trap was used, MS2 were acquired using turbo scans covering a mass range of 150–1250 *m/z*. The Orbitrap methods collected MS2 spectra at 15,000 resolution covering a range of 150–2000 *m/z*.

#### Replicate injections of human plasma digest

To assess technical precision, 3 *μ*L of the same vial of 100% human plasma digest was injected in five separate replicates. Each replicate consisted of four total injections, one injection using each of the four methods, run in a random order.

#### Matrix-matched calibration curve

The calibration curve was created by acquiring three replicates of each dilution point (**Table S1**) in both the linear ion trap and Orbitrap both using only the 1.7 Th precursor isolation width. Data were acquired starting with the lowest dilution point, running one injection each in the linear ion trap and Orbitrap, then moving to the next lowest dilution point until the full curve was run. Following a quality control run, the process was repeated for a total of 3 replicates. The order of linear ion trap and Orbitrap acquisition was randomly assigned for each replicate of each dilution point.

#### Data availability

The raw MS files, EncyclopeDIA .elib file, and all Skyline documents are available have been deposited in the ProteomeXchange consortium^12^ (identifier PXD023334) via Panorama Public^13^ (https://panoramaweb.org/qLITvsqOT.url).

### Data processing and analysis

#### Library creation and refinement

A chromatogram library was created by searching gas-phase fractionation DIA data against the Pan-Human library^14^ in EncyclopeDIA (v 0.9.0),^15^ using a method that has been described in detail previously.^11^ The chromatogram library was loaded into Skyline (v 20.2.1.315)^16,17^ along with the 8-*m/z* DIA data collected for this experiment demultiplexed and converted to mzML format in MSConvert (Proteowizard, V 3.0.20239).^18^ From these data, the five most abundant peptides were selected for each protein, and the remaining peptides were manually filtered for a high dotP, high retention time correlation with the chromatogram library, and minimal interference. Preference was given to peptides from plasma proteins that have been previously identified for their role in monitoring human wellness (https://panoramaweb.org/Passport).^19^ The isolation list of all 432 peptides plus 14 PRTC peptides was scheduled with a 5 minute retention time window (**Figure S1**). As PRM data were acquired, results were loaded into Skyline and used to adjust scheduling windows as needed.

#### Data parsing

Raw files from the PRM run were imported into Skyline Daily (v 20.2.1.315)^16,17^ for peak detection, integration, and visualization. Runs using the linear ion trap were integrated using a tolerance of +/− 0.7 *m/z*, while the Orbitrap was processed with a tolerance of 20 ppm. Peak picking was manually refined for all data files, and area under the curve abundance was obtained for each peptide. For the replicate injection experiment, all peptide abundances were normalized to the median signal of the run. For the matrix-matched dilution curve runs, peptide abundances were normalized to the global standards (PRTC peptides). Thirty-seven peptides that were observed to have truncated peaks in one or more of the matrix-matched dilution curve runs or to be present in the chicken digest were removed from this analysis. Area under the curve data were exported from Skyline, and analyzed in R (v 3.6.1) and Python (v 3.8.2). Limits of quantitation (LOQs) were calculated for each peptide in each analyzer from the dilution curve data using the method described by Pino et al. (2020).^20^ The code used for analysis and figure generation is available on Github (https://github.com/uw-maccosslab/iontrap_vs_orbitrap).

#### Transition refinement to optimize limit of quantitation

A list of areas under the curve for all singly charged b- and y- ions beyond the second residue were exported from Skyline. For each peptide and each mass analyzer, the optimal set of transitions was determined independently as follows. First, the LOQ was calculated for all transitions separately using the method developed by Pino et al.^20^ The LOQ was then calculated for the summed area of the four transitions with the lowest LOQ, which constituted the initial accepted transition set. Then, the LOQ was calculated for the accepted transition set and each of the remaining transitions. Each additional transition that lowered the LOQ was added to the accepted transition set. Finally, the refined LOQ was recalculated with the full set of transitions.

## Results and Discussion

### Creation of the PRM assay

To compare mass analyzers, we chose plasma as a sample matrix because it is clinically relevant and its large dynamic range^21,22^ is well suited for testing the limits of targeted peptide quantitation. To develop a robust PRM assay, we started with a DIA-based chromatogram library consisting of 3084 peptides mapping to 358 proteins. This library was developed using gas-phase fractionated DIA data that was searched against the Pan-Human library^14^ in EncyclopeDIA.^15^ The resulting chromatogram library contained information on the identity, fragmentation spectra, and retention times of all detectable peptides. This original library was created using a different instrument, liquid chromatography, and sample than those used in this PRM experiment. Therefore, the library needed to be refined to create a scheduled targeted method that could be acquired with a sufficient number of points across each chromatographic peak.

To generate a targeted assay from the existing chromatogram library, a DIA run was collected using staggered 8-*m/z* windows with the Orbitrap mass analyzer. The resulting data were used to determine accurate retention times for all peptides. Based on these data, the original peptide library was filtered for peptides that were 1) confidently seen in this sample, 2) biologically relevant (based on a previous desire to focus on proteins useful for assessing general wellness), and 3) came from proteins spanning a wide dynamic range (**Figure S2**). To fulfill these criteria, the five peptides per protein with the most intense peak areas from the product ion data were selected, and the list was further refined to ensure all peptides had high dotp with the library chromatograms. Additionally, peptides were preferentially selected if they came from proteins that have been used in other MS-based targeted plasma assays.^19^ The final assay consisted of 432 peptides derived from 128 protein coding genes such that upon scheduling no more than 75 peptides would be measured at any given time using a five-minute retention time window (**Figure S1**).

The primary consideration in PRM assay development is limiting the number of targets monitored at one time to maintain a short enough cycle time to sample at least eight points across each peak without sacrificing sensitivity. In the Orbitrap, the length of time required for spectral acquisition does not depend on the range of mass-to-charge, because of the multiplexing property of Fourier transform-based analysis. As a scanning instrument, the linear ion trap pays a time penalty proportional to the range of mass-to-charge acquired, however the total scan time for the range 150–1250 *m/z* is only 8.8 ms with the 125 kTh/s scan speed. On both the Orbitrap and linear ion trap, the analysis takes place while the collision cell is being filled with ions for the next spectrum, so the cost of mass analysis to the overall duty cycle is quite low. The time allotted for ions to accumulate prior to spectral acquisition is typically the limiting factor in cycle time and is crucial for maintaining sensitivity. Critically, both analyzers maintain a fundamental advantage over triple-quadrupoles because all transitions are collected within a single MS/MS spectrum and therefore can be refined after data acquisition. In contrast, each additional transition measured costs an additional dwell/acquisition time in SRM. This benefit of PRM eliminates the need to perform time consuming experiments to select interference-free transitions prior to acquisition, as is the case when transferring accurate mass data to triple-quadrupole assays.

Although high-resolution mass analyzers have traditionally been favored for PRM, linear ion traps may be ideally suited for PRM experiments due to their speed, sensitivity, and ability to scan whole mass ranges without significantly raising the cycle time.^23^ Still, the speed and sensitivity of linear ion traps come at a cost to the resolution, which could present challenges for PRM experiments. First, it may be difficult to select the correct peak for a peptide over the course of a run, there can be multiple peptides with similar precursor masses and shared product ions which generate multiple possible peaks (**Figure S3**). By utilizing a chromatogram library with retention times and fragmentation patterns calibrated on a high resolution instrument, it is possible to build an assay without using synthetic standards to ensure accuracy. A second potential problem with unit resolution is the increased potential for chemical interference, leading to inaccurate quantitation. However, we argue that this problem can be addressed simply by selecting interference-free fragment ions. Furthermore, it is possible that the increased speed and sensitivity of a q-LIT could make it better suited for PRM assays than its high resolution counterparts.

### Evaluation of technical precision

To assess the impact of precursor and product ion selectivity on technical precision, the same sample of human plasma digest was injected and measured using both the linear ion trap and Orbitrap each with two different precursor isolation widths. By collecting multiple replicates with the same sample on the same instrument, it was possible to compare the technical variance of the mass analyzers directly. Based on five replicate injections, the percent coefficient of variation (% CV) was determined for each peptide in each method (**Figure 2A**). The Orbitrap had lower technical variation than the linear ion trap for most peptides (**Figure 2B**), but the differences in variation would likely make a minor difference in practice. At the 1.7 Th precursor isolation width, the median CV in the linear ion trap of 5.8% was slightly higher than the Orbitrap’s 4.1%. Notably, both median CV values are more than adequate for quantitative experiments and at the same 1.7 Th precursor isolation width, the linear ion trap had a higher proportion of peptides, 98.3%, than the Orbitrap, 97.6%, below an arbitrarily selected 20% CV cutoff. The 20% CV cutoff is frequently used as a threshold for quantifiable peptides, and is an important factor in comparing these performances. While the Orbitrap maintains slightly better median precision than the linear ion trap, the practical differences in precision between using these two analyzers are small.

**Figure 2.**
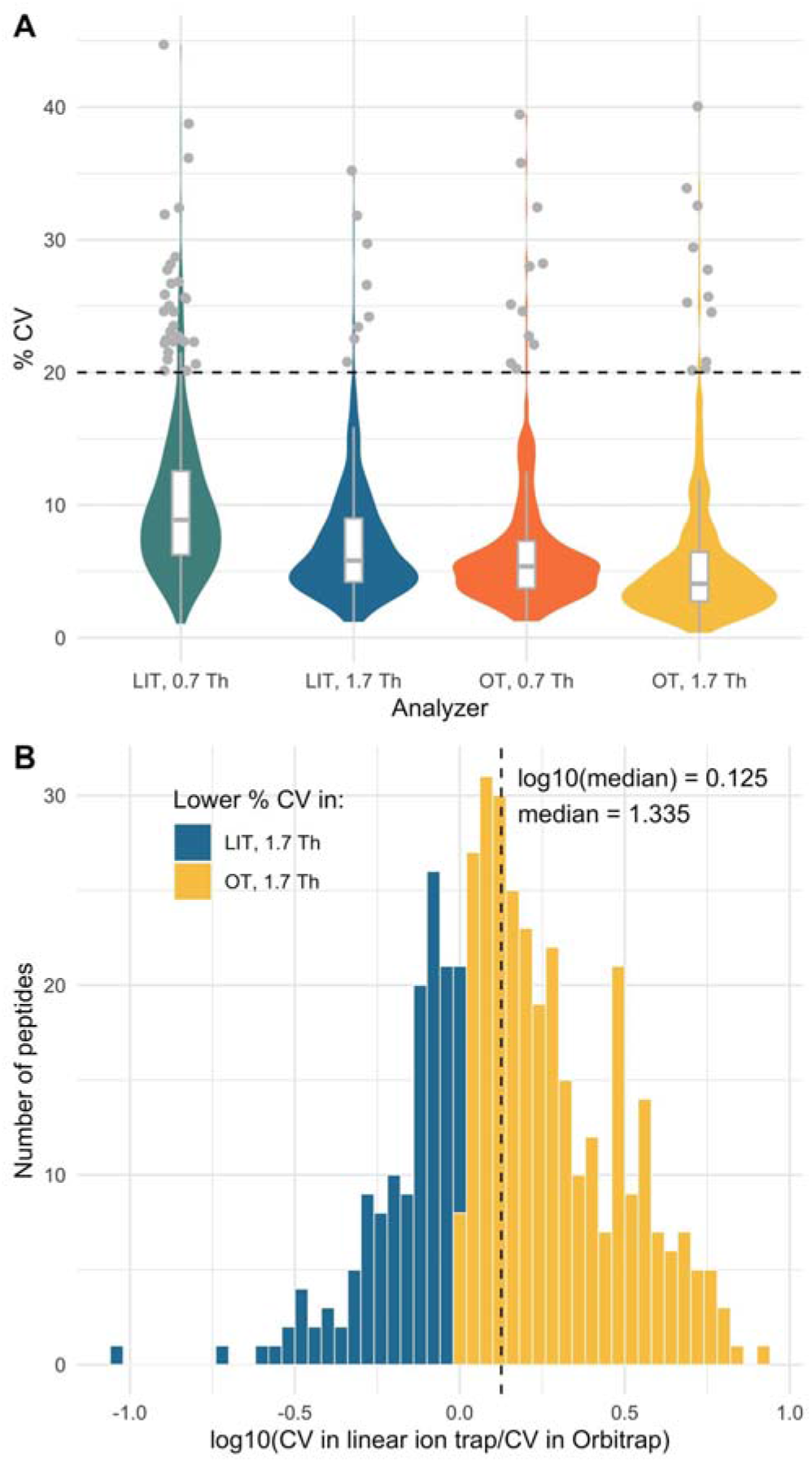
Comparison of coefficients of variation between PRM data acquired with the orbitrap (OT) and the linear ion trap (LIT). CV’s for each peptide in the assay when monitored with both mass analyzers at two precursor isolation widths (0.7 and 1.7 Th) (A). The log ratio of every peptide CV in linear ion trap and Orbitrap, where a positive log ratio indicates a better CV in the Orbitrap than linear ion trap (B).

One factor that can lead to high technical variation is the presence of interfering species. Interference can occur when an interfering precursor ion is co-isolated with the target precursor ion and generates product ions that fall within the integration boundaries of the target product ions, creating a shoulder in one or more product ion chromatograms (**Figure S4A and B**). Between runs, slight changes in chromatography can shift the relative position of the target peptide and interfering species, causing significant changes in quantitation (**Figure S4C**). Chemical interference of this nature can be influenced by both the precursor and the product ion selectivity.

Precursor ion selectivity is partially determined by precursor isolation width. If a narrower isolation window is used, potentially interfering species may not be co-isolated at all, eliminating that interference. However, reducing the isolation window comes at a cost to sensitivity as the narrow quadrupole transmits fewer ions. Therefore, quadrupole isolation width should be adjusted to strike a balance between sensitivity and selectivity. For both the Orbitrap and the linear ion trap, the 0.7 Th precursor isolation window tended to have higher technical variance because the traps tended to fill with fewer ions (**Figure S5**). When there are fewer ions to measure the relative variance tends to increase^24^ which explains the increased precisions with a wider precursor isolation window. Both analyzers performed better with the 1.7 Th isolation width, so those methods were used for the remaining experiments and analyses.

Similarly, product ion selectivity is related to the MS/MS resolution. In a lower resolution spectrum, there is a larger chance that another analyte will have an indistinguishable *m/z* at a specific retention time and is therefore more likely to have interference. This difference is reflected in the necessary integration boundaries for each analyzer: in the unit resolution linear ion trap, we must integrate the area +/− 0.7 *m/z* around the theoretical peak, whereas the Orbitrap only requires integration boundaries of 20 ppm. Thus, the unit resolution linear ion trap has the potential to have more interference than the Orbitrap, but with the careful selection of transitions, this problem may be minimized (**Figure S6**).

The fear of increased variance due to increased interference has been a major reason why the proteomics community has largely trended towards using high resolution mass analyzers for PRM. Indeed, the transition set used in this experiment is predicted to have more interference for the linear ion trap than the Orbitrap (**Figure S7**), although this prediction does not account for separation of potential interfering species across time. In fact, our results suggest that chemical interference of this nature was minimal, even in the linear ion trap. Thus, the Orbitrap’s superior performance can be attributed to several factors, including the fact that it had a higher AGC target and tended to measure more ions per spectrum than the linear ion trap (**Figure S5**), a factor that has been demonstrated to improve precision.^25–27^ Still, the Orbitrap tended to hit its maximum inject time in most cases (**Figure S5C**) and was below its target fill capacity, which is likely why the difference in precision between the analyzers is not more pronounced.

### Assessment of sensitivity with matrix-matched calibration curves

To assess the sensitivity of each analyzer, we determined the lower limit of quantitation for each peptide using a matrix-matched calibration curve.^20^ To determine the LOQ for all peptides in the assay, an 11-point dilution curve was prepared by diluting human plasma in chicken serum (**Table S1**). This matrix-matched method allows for the dilution of analytes without significantly changing the matrix composition.^20,28^ To reduce run-order bias, the complete dilution curve was run with each method from low to high concentration, randomizing the order of analyzers within each point of each replicate, and the process was repeated for a total of three replicates. Data were collected by scheduled PRM using either the Orbitrap or LIT with a 1.7 Th precursor isolation window to acquire the product ion spectra. The PRM data were imported into Skyline where the area under the curve was assessed for each peptide. A report was generated with the peak areas and analyzed using a script modified in house based on the method developed by Pino et al.^20^ This method uses a bilinear fit to assess the turning point in the calibration curve and then uses a bootstrapping method to find the lowest concentration where the CV was below 20%. This powerful method has many benefits over alternative methods, including that it is able to estimate LOQs between dilution points, instead of simply finding the lowest measured quantitative point.^20^ The ratio of the LOQ for any given peptide in each mass analyzer could be used to compare these results directly.

Peptides tended to have lower limits of quantitation in the linear ion trap compared to the Orbitrap (**Figure 3A**). To obtain a more complete understanding of analyzer performance, the LOQ for each peptide was compared individually by dividing the LOQ in the linear ion trap by the LOQ in the Orbitrap. A ratio less than one indicates that the linear ion trap has a lower (better) LOQ, whereas the reverse is true for a ratio greater than one. The distribution of these ratios skews in favor of the linear ion trap, with a median ratio of 0.858 (**Figure 3B**). This ratio indicates that LOQs in the linear ion trap were generally lower than those in the Orbitrap, suggesting the linear ion trap more than compensates for the greater potential for interference. The sensitivity of the analyzer can be just as critical as precision in evaluating performance— more sensitive mass analyzers require less analyte to generate meaningful signal. We use the lower limit of quantitation, or the minimum point on a dilution curve where a change in signal reflects a change in quantity, to assess sensitivity quantitatively.^29^ Based on this analysis, we determined that the linear ion trap tends to be more sensitive than the Orbitrap for this PRM assay.

**Figure 3.**
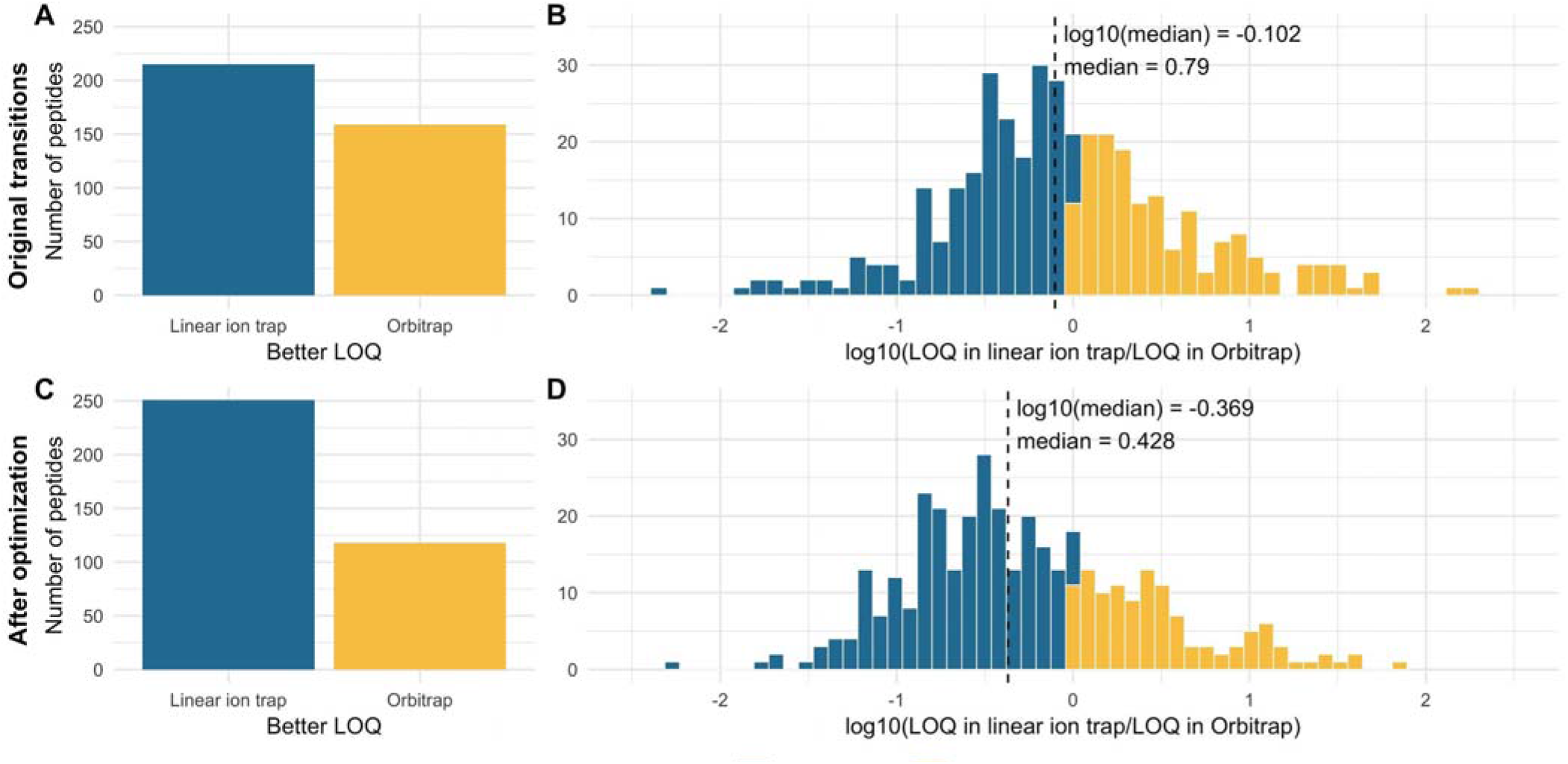
LOQ’s before (A&B) and after (C&D) transition refinement. A and C show the raw number of peptides with better LOQs in each analyzer. B and D show the log ratio of LOQs, with a log ratio greater than 0 meaning the Orbitrap had a better LOQ than the linear ion trap. Dashed line indicates the median ratio. Before and after optimization, 21 and 26 peptides respectively had limits of quantitation outside the dilution curve in both analyzers and had identical log ratios (not shown).

Theoretically, the sensitivity of each analyzer is limited by the number of ions it requires to produce a quantitative signal. In a targeted experiment on a trapping instrument, the number of ions in each trap is determined by the ion current, which is inherent to the sample and injection setup, and the maximum inject time, a parameter set based on the number of concurrent targets and the desired cycle time. Highly abundant peptides may fill the trap with the desired number of ions in less time, while less abundant peptides may take much longer to fill the trap. In cases where the maximum inject time is too short to fill the trap with the minimum number of ions to obtain sufficient signal, the variation in the measured peptide abundance may be too large to be considered “quantitative”. Thus, one crucial difference between the linear ion trap and Orbitrap is the number of ions needed to produce a detectable signal. The linear ion trap uses electron multiplier based detection, with gain sufficient to detect single ions.^23^ In contrast, Makarov and Denisov demonstrated Orbitrap detection limits of 20+ charged single myoglobin ions with signal-to-noise 3.5 using 0.76 s transients.^30^ With the 0.032 s transient, 15k resolution setting used in our experiments, the minimum detectable number of ions should be ~5x higher, because signal-to-noise ratio varies as the square root of transient length. Therefore, with these settings the Orbitrap needs 1–2 orders of magnitude more ions to create a detectable signal than the linear ion trap. This greater theoretical sensitivity explains why the linear ion trap performs better in this experiment despite the increased potential for interference.

### Automated transition refinement

The original transition set was selected to minimize interference in the chromatogram library generated from non-diluted plasma on an Orbitrap and further refinement is required to optimize the transition set for this experiment. To this end, transitions were automatically selected to produce the lowest limit of quantitation (**Figure S8**). For both analyzers, the refined transitions yielded better LOQs for most peptides, and unsurprisingly this change was more pronounced in the linear ion trap than in the Orbitrap (**Figure 4**). Consequently, after this transition refinement, the linear ion trap had even better LOQs relative to the Orbitrap than before (**Figure 3C**). The median ratio of LOQs in the linear ion trap compared to the Orbitrap went from 0.790 to 0.428, indicating that the LOQs in the linear ion trap had a larger net improvement, and the linear ion trap outperformed the Orbitrap by an even more significant margin (**Figure 3D**).

**Figure 4.**
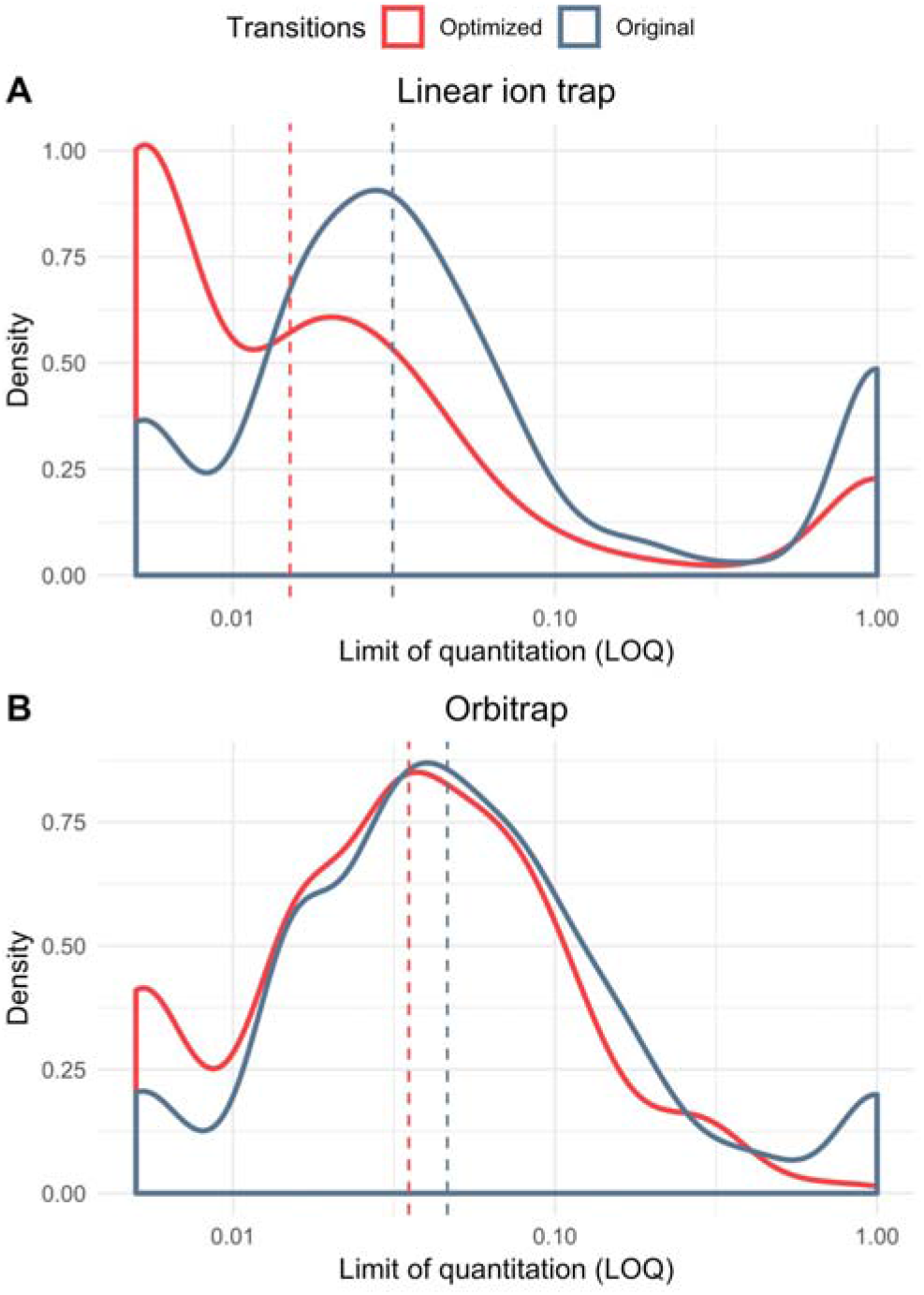
Probability density function (PDF) of limit of quantitation before and after transition refinement in (A) linear ion trap and (B) Orbitrap. The x-axis corresponds to the lower limit of quantitation in log scale with the median of each transition set indicated by the appropriately colored dashed line. A lower x-value indicates a lower LOQ, meaning better sensitivity. Both analyzers have better LOQs after transition selection, but the linear ion trap improves more as a result of optimization.

The observation that transition refinement had a larger impact on the linear ion trap than the Orbitrap may be a result of the assay construction. Because this assay was based on a chromatogram library constructed using interference-free transitions determined using a different sample measured on an Orbitrap, the transitions were never optimized for these specific samples and instrument configuration. The library was generated with an undiluted human plasma digest and in the case of a similar undiluted sample such as the one used to determine precision, the chosen peptides had minimal interference regardless of the mass analyzer. However, as the dilutions with the matched matrix increased, interference increased as the transitions were not optimized for these lower dilution points. Additionally, the original transitions selected to be interference-free were selected based on Orbitrap data, and it is likely that many of these transitions may not be optimal in the linear ion trap. Based on the improved mass accuracy and resolution of the Orbitrap, we predicted interference at different precursor/product ion specificities and the original peak picking should have been better for the high resolution Orbitrap than the unit resolution linear ion trap (**Figure S7**). Furthermore, peptides that had a lower LOQ in the linear ion trap qualitatively lacked signal at low dilution points using the Orbitrap (**Figure S9A**). Conversely, peptides with a lower LOQ in the Orbitrap subjectively had interference at lower dilution points in the linear ion trap (**Figure S9B**). These subjective observations are supported by the fact that the linear ion trap quantitative data improves significantly more than the Orbitrap after transition refinement (**Figure 4**).

The ability to refine transitions based on empirical results is a major benefit of using an instrument that acquires full MS2 spectra over a triple quadrupole where transitions must be selected prior to analysis. It is possible to reduce interference post-acquisition simply by selecting a transition set to minimize interference. In both mass analyzers, almost all peptides are predicted to have at least four interference-free transitions (**Figure S6**), although the correct transitions may not have been selected in the original assay. When selecting transitions, it is preferable to select as many interference-free transitions as possible to increase the number of ions measured thereby increasing precision.^25–27^ Thus, the transition refinement method presented here selects at least four transitions because at this threshold, there is very little difference between analyzers in theoretical interference (**Figure S6**). This automated transition refinement heuristic is a simple method to leverage the full power of PRM to accurately quantify peptides.

## Conclusions

Here, we show that the power of PRM lies in its ease of use, sensitivity, and flexibility in selecting transitions post-acquisition—rather than high mass accuracy. We report that unit resolution PRM performs at least as well as high resolution PRM in this targeted plasma assay by demonstrating that of the total set of 432 assayed peptides, over 97% could be quantified precisely (<20% coefficient of variation) with both analyzers, and the linear ion trap produced better lower limits of quantitation compared to the Orbitrap for 62% of all peptides. High resolution spectra and mass accuracy are not a requisite for selectivity, especially when peptide transitions can be refined post-acquisition. While not addressed here, in the rare event where unit resolution tandem mass spectrometry might provide insufficient selectivity, the linear ion trap has the capability to perform multistage MSn, whereas the Orbitrap is limited to MS2 without the LIT. Overall, the superior speed and sensitivity of the linear ion trap on the Orbitrap Eclipse make it better suited for targeted proteomics than the use of an orbitrap mass analyzer.

## Notes

The authors declare the following competing financial interest(s): The MacCoss Lab at the University of Washington has a sponsored research agreement with Thermo Fisher Scientific, the manufacturer of the instrumentation used in this research. M.J.M. is a paid consultant for Thermo Fisher Scientific. P.M.R. is an employee of Thermo Fisher Scientific, the manufacturer of the instrumentation used in this research.

## Acknowledgements

This work is supported in part by National Institutes of Health Grants P41 GM103533, U01 DK121289, and U19 AG065156.

## Supporting Information

**Supplemental File S1**: Supplemental figures and tables referenced in the text.

## Notes

https://panoramaweb.org/qLITvsqOT.url

https://github.com/uw-maccosslab/iontrap_vs_orbitrap

