## Supplemental File S1 for "Comparison of unit resolution versus high-resolution accurate mass for parallel reaction monitoring"

### Supplement

| Label | % Human | % Chicken |
|---|---|---|
| A | 0 | 100 |
| B | 0.5 | 99.5 |
| C | 1 | 99 |
| D | 3 | 97 |
| E | 5 | 95 |
| F | 7 | 93 |
| G | 10 | 90 |
| H | 30 | 70 |
| I | 50 | 50 |
| J | 70 | 30 |
| K | 100 | 0 |

**Table S1**. List of points on the dilution curve. Dilution curve was made up of 11 points, with more points at low concentration to get more accurate lower limits of quantitation.

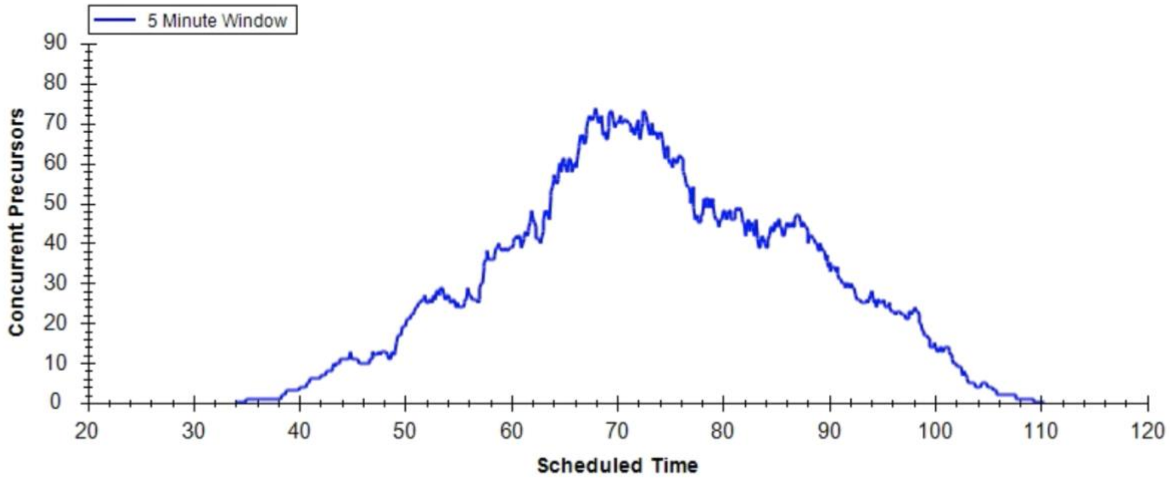


**Figure S1**. Number of precursors monitored at any given time throughout the assay. Scheduling was based on measured retention time with a 5-minute window. Retention times were updated throughout the experiment to account for small changes in chromatography.

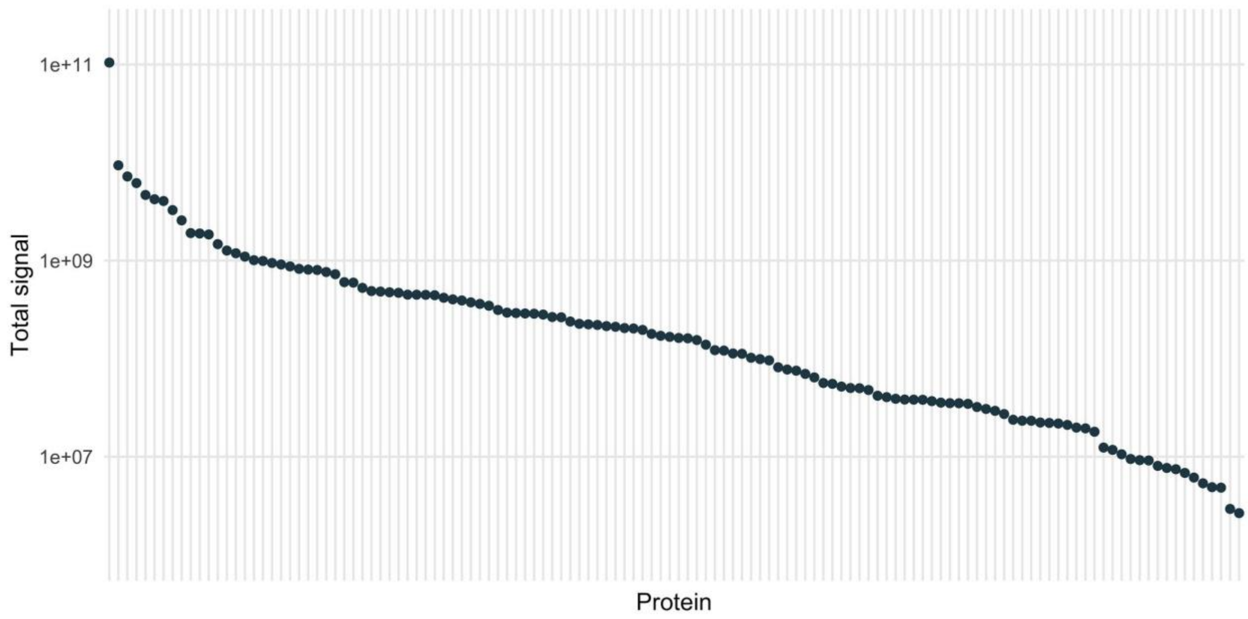


**Figure S2.** Plot of dynamic range of all proteins in the assay. Each point represents the summed area under the curve abundance of all peptides mapping to that protein. The total signal ranges from $1.0\times10^{11}$ for serum albumin down to $2.6\times10^{6}$ for vitamin K-dependent protein C.

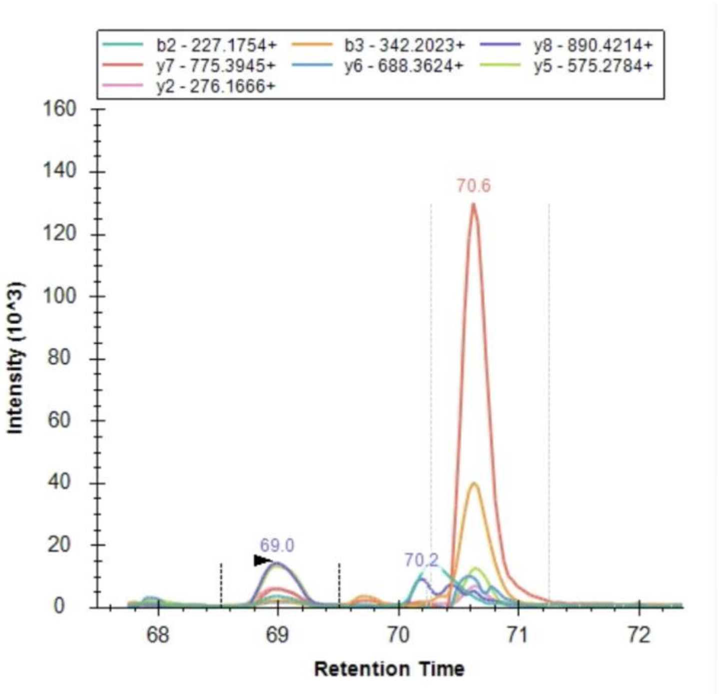


**Figure S3.** Extracted ion chromatograms for all product ions in a 5-minute window for one precursor collected with linear ion trap. Actual peak at 69,0 minutes could be determined based on retention time and accurate product ion fragmentation patterns. Without retention time information, this determination could be difficult.

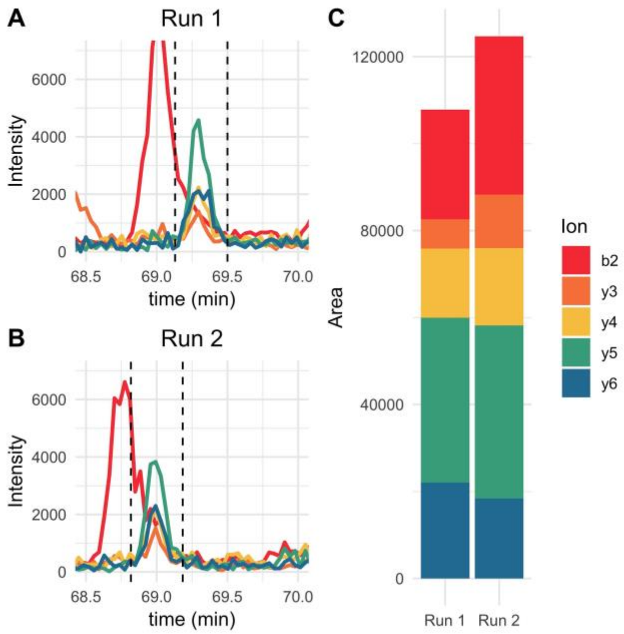


**Figure S4.** Example chromatograms of a peptide with interference. Chromatograms show significant interference impacting the b2 ion (A&B). The variance of the integrated peak area for this peptide is mostly influenced by the b2 ion getting integrated slightly differently in both runs (C).

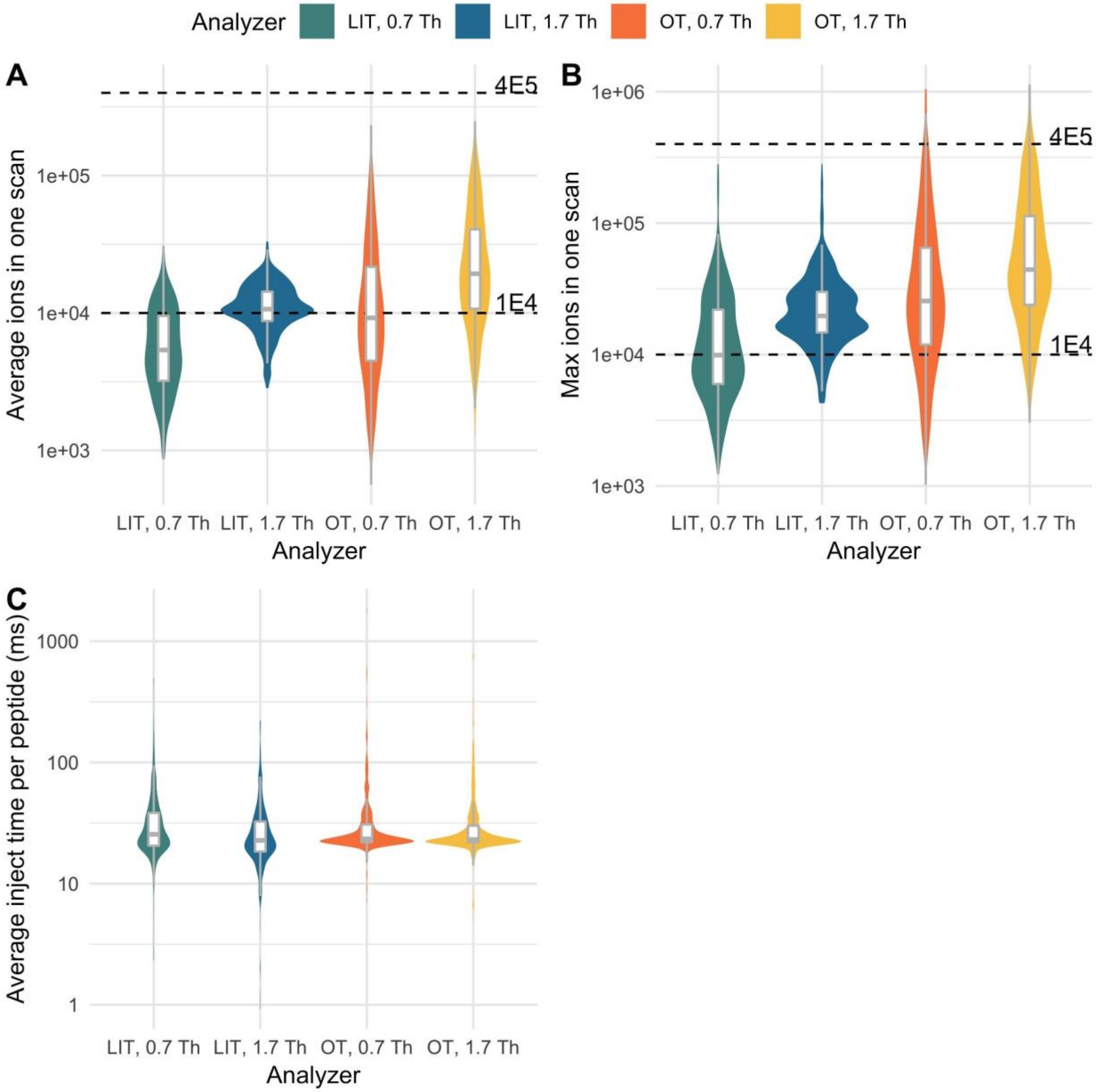


**Figure S5**. Ion fills and inject times for each analyzer. Plots display the average number of ions per scan (A) or maximum number of ions in a scan (B) for all peptides in each mass analyzer. The number of ions per scan was calculated by multiplying the total ion current by the inject time for that scan and dividing by 1000. For the Orbitrap, this value was then multiplied by 0.0917, an instrument and transient length specific conversion factor obtained by measuring the ratio of ion trap peak intensity to Orbitrap peak intensity for a series of calibration ions. When a narrow quadrupole isolation width is used, the mean and max number of ions gets pretty low, which would explain the higher variance at these isolation widths. Another metric to look at fill is the average inject time per spectrum in each of the analyzers (C). Orbitrap has a narrower distribution of inject times, likely because the maximum inject time was hit more frequently.

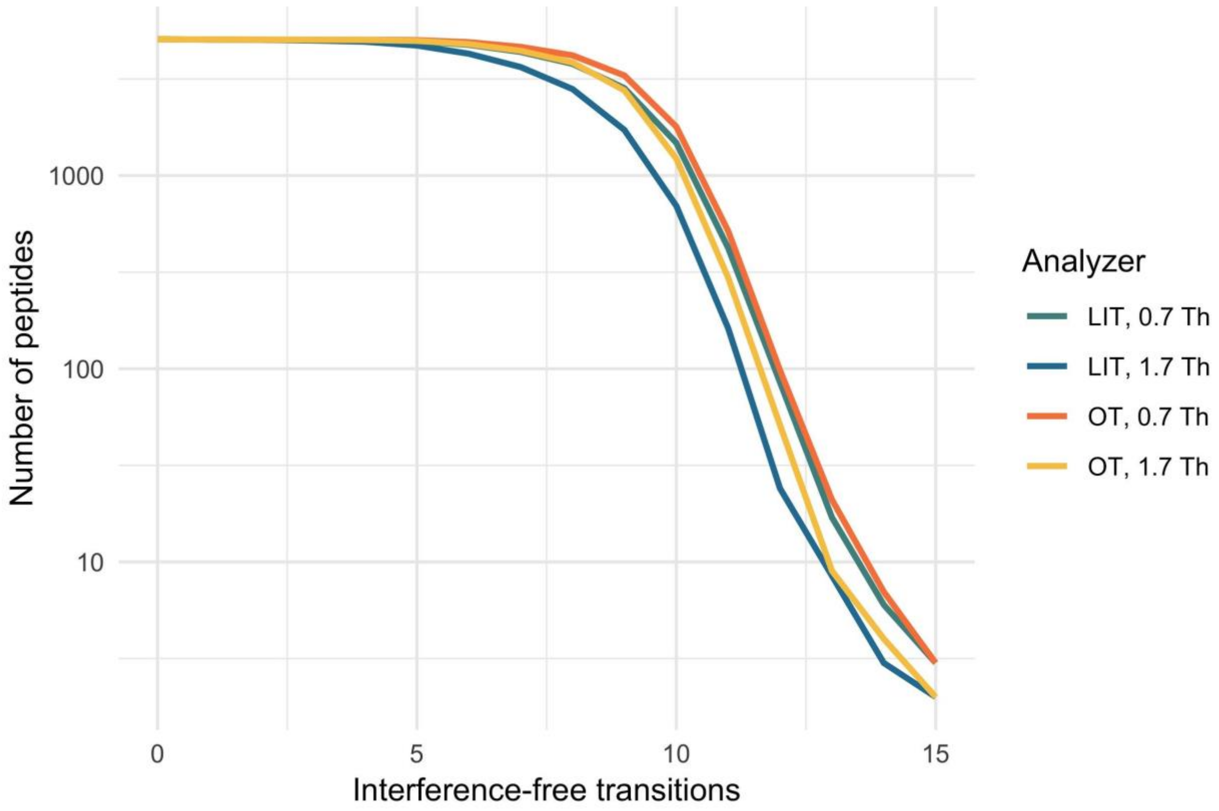


**Figure S6**. Predicted number of interference-free transitions for all peptides in the plasma chromatogram library. The y-axis represents the number of peptides with *x* transitions or more. Interference-free transitions are defined as precursor/ y-ion pairs that do not overlap within the appropriate *m/z* tolerance with b- or y- ions generated by precursors within the specified isolation window. At 1.7 Th, the linear ion trap is more susceptible to interference in general, but there is little loss if four or fewer transitions are selected.

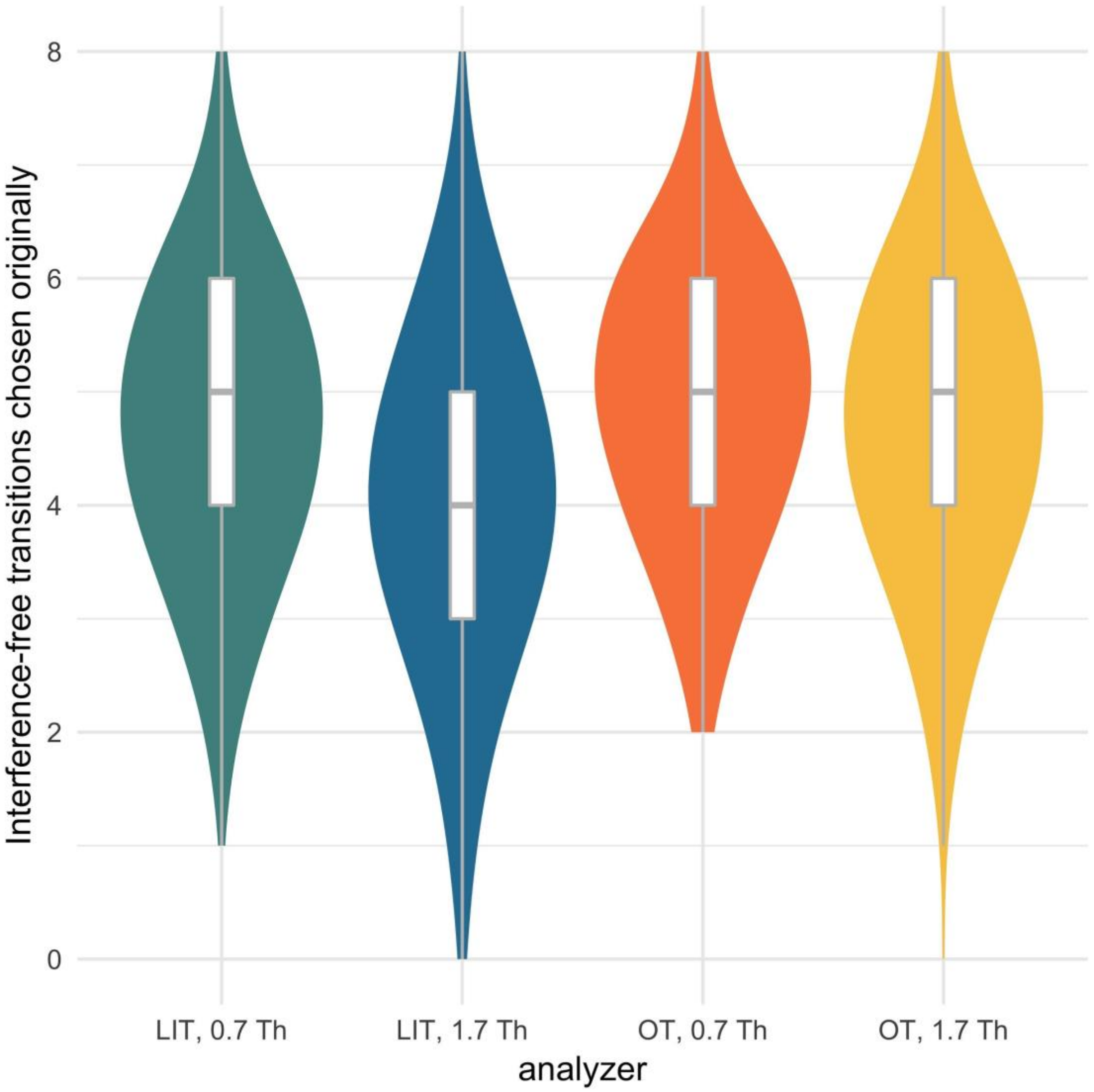


**Figure S7**. Number of theoretically interference-free transitions for each experiment. The original transition set has more interference for linear ion trap with the parameters used in matrix-matched experiment (LIT, 1.7 Th compared to OT, 1.7 Th). However, even in this set, most peptides have at least four interference-free transitions before refinement.

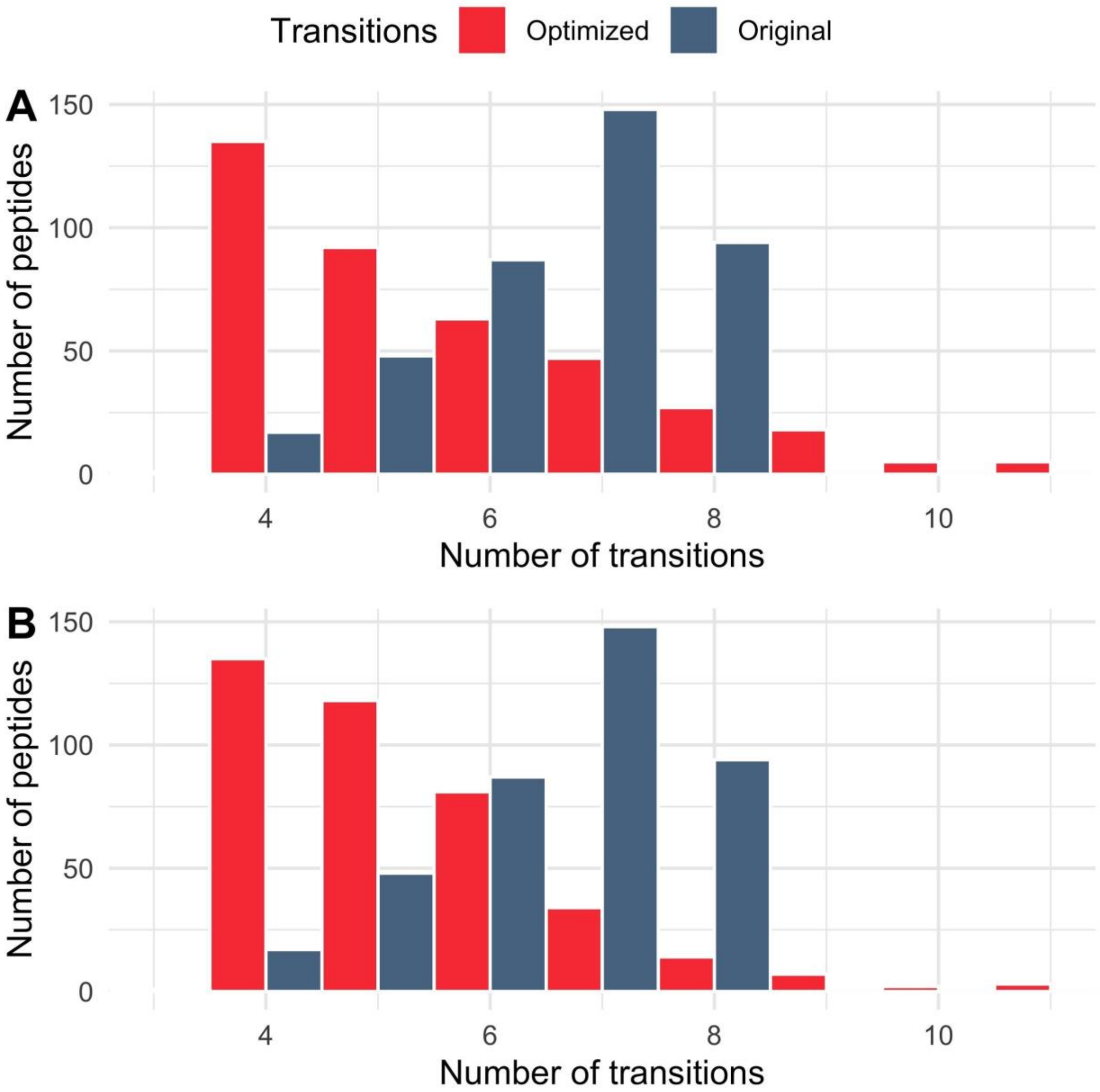


**Figure S8**. Number of transitions selected before and after refinement in (A) linear ion trap and (B) Orbitrap. Before refinement, there were generally more transitions in both analyzers, indicating that this method of manually refining transitions is able to remove interference that crops up at low dilution points. The mode of the optimized distribution is also the minimum number of possible transitions, which is probably due to the fact that many transitions have interference when the signal is low.

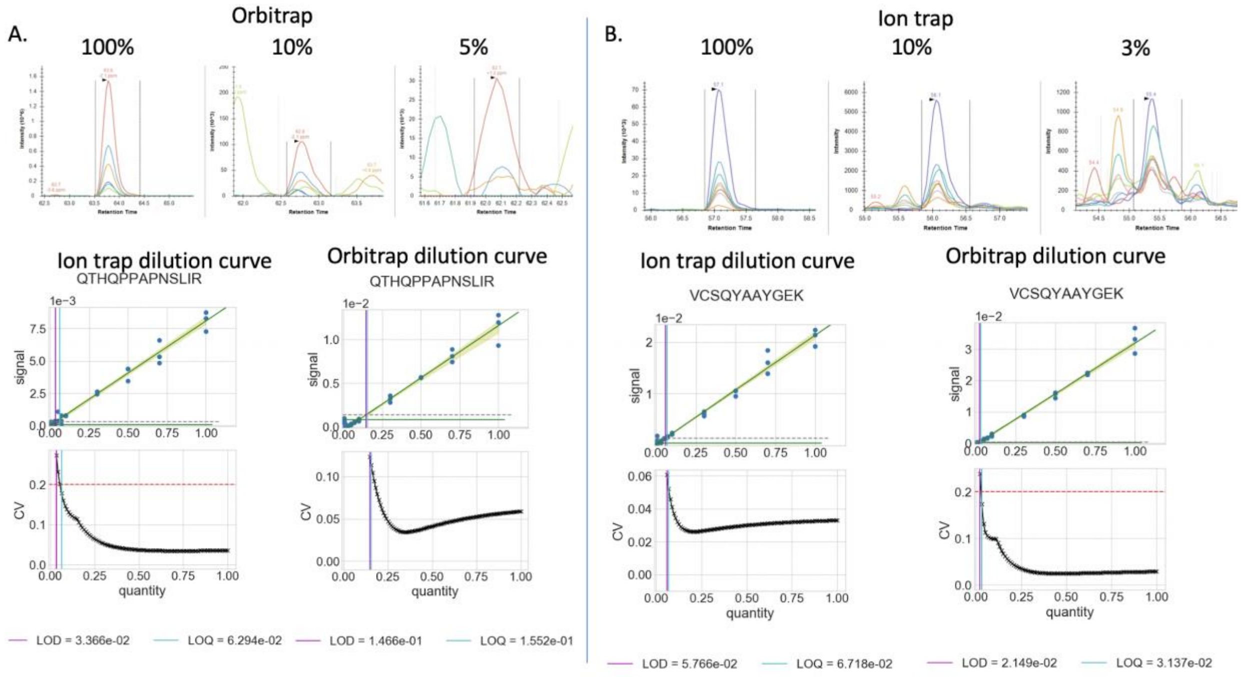


**Figure S9.** Examples of peptides with better LOQ in linear ion trap (A) and Orbitrap (B). Top panels in each figure show what happens to signal in the analyzer with the worse LOQ. In the case of Orbitrap (A), signal fades away at low points. On the other hand, interference becomes more apparent at lower dilutions when the linear ion trap performs worse (B). The bottom panel shows dilution curves in both analyzers for each peptide.
